# An inverted Caveolin-1 topology defines a novel exosome secreted from prostate cancer cells

**DOI:** 10.1101/2020.03.19.999441

**Authors:** Nicholas Ariotti, Yeping Wu, Satomi Okano, Yann Gambin, Jordan Follett, James Rae, Charles Ferguson, Rohan D. Teasdale, Kirill Alexandrov, Frederic A. Meunier, Michelle M. Hill, Robert G. Parton

## Abstract

Caveolin-1 (Cav1) expression and secretion is associated with prostate cancer (PCa) disease progression but the mechanisms underpinning Cav1 release remain poorly understood. Numerous studies have shown Cav1 can be secreted within exosome-like vesicles, but antibody-mediated neutralization can mitigate PCa progression; this is suggestive of an inverted (non-exosomal) Cav1 topology. Here we show that Cav1 can be secreted from specific PCa types in an inverted vesicle-associated form consistent with the features of bioactive Cav1 secretion. Characterization of the isolated vesicles by electron microscopy, single molecule fluorescent microscopy and proteomics reveals they represent a novel class of exosomes ∼40 nm in diameter containing ∼50-60 copies of Cav1 and strikingly, are released via a non-canonical secretory autophagy pathway. This study provides novel insights into a mechanism whereby Cav1 translocates from a normal plasma membrane distribution to an inverted secreted form implicated in PCa disease progression.

## INTRODUCTION

Caveolae are a characteristic feature of the plasma membrane (PM) of many mammalian cell types^1^. Caveolin-1 (Cav1) is the major non-muscle isoform of the caveolin family and is essential for caveola formation^2^. Cav1 is synthesised at the endoplasmic reticulum and exported through the Golgi complex to form caveolae at the cell surface^3^. PM-associated caveolae can be internalised and fuse with early endosomes before recycling back to the surface without disassembly^4, 5^. A peripheral membrane protein, termed Cavin1 or Polymerase Transcript Release Factor (PTRF) is also required to stabilise caveolae on the PM^6, 7^. The loss of Cavin1 results in a switch from a largely stable pool of Cav1 at the PM to a rapidly internalised pool with increased endocytic recycling^6, 8, 9^. It has been postulated that the ratio of Cav1 expression to Cavin1 expression is a key regulator of Cav1 dynamics as this ratio is tightly regulated in different cell types and tissues^8^.

Cav1 over-expression correlates with advanced and metastatic prostate cancer (PCa)^10–12^ in the absence of Cavin1 expression^11^. A loss of Cav1 expression in the TRAMP transgenic mouse PCa model resulted in a dramatic reduction in PCa growth and metastasis^13^. Moreover, PCa-associated Cav1 exists in dynamic non-caveolar membrane domains as observed in the aggressive PC3 cell line, which expresses Cav1 but not Cavin1^6^. Cavin1 expression is sufficient to attenuate Cav1-associated PCa disease progression in an orthotopic xenograft mouse model^11^. In addition to reducing anchorage-independent growth and migration, Cavin1 expression altered the secretion of Cav1 ^14, 15^ as well as the tumour microenvironment^16^.

Cav1 is a potential paracrine factor modulated by Cavin1 expression. PCa cell-secreted Cav1 has been shown to enhance cancer cell survival, to stimulate PCa cell proliferation^17^, and to have pro-angiogenic effects^18^. The effect of Cav1 is clearly mediated by secreted caveolin as it can be recapitulated with medium from LNCaP cells heterologously expressing Cav1^19^, and can be inhibited by antibodies to Cav1 in cultured cell systems and mouse models^12, 17^. Moreover, serum Cav1 is elevated in PCa patients compared with control men or men with benign prostatic hyperplasia^20^, and is a potential prognostic marker for PCa recurrence after prostatectomy^21^. Elucidating the various mechanism(s) by which caveolin is secreted is therefore critical to understand Cav1’s function in PCa progression.

Considerable evidence from the literature shows that Cav1 can associate with exosomes^14, 19, 22, 23^ and is highly secreted within exosomes derived from PC3 cells^14, 24, 25^. Exosomes are formed in the endocytic pathway and comprise a subset of intralumenal vesicles (ILVs) within the multivesicular bodies (MVBs)^26^. The ∼100-200 nm vesicles are released from the cell when MVBs fuse with the cell surface^27^. As the exosome is a membrane-bound vesicle, integral membrane proteins are released from the cell within a membrane bilayer in an energetically favourable lipid environment ^28^. However, an exosome release model for secretion of bioactive caveolin is consistent with some, but not all, published data. First, exosomes are typically sedimented from the medium at 100,000 x *g*, however several studies have reported that secreted Cav1 is not pelleted under these conditions^12, 19^. Second, antibodies against caveolin have been shown to inhibit the effect of secreted caveolin^12, 17, 19^: a released exosome would contain all exposed caveolin epitopes masked within the lumen of the vesicle, which is topologically equivalent to the cytoplasm. Third, caveolin has been demonstrated to be secreted through other non-classical means in numerous studies via an undefined mechanism^29, 30^.

In this study we have used a variety of biochemical assays and microscopy-based imaging techniques to interrogate the route of caveolin release from prostate cancer cell lines. We confirm Cav1 is released within conventional exosomes from PC3 cells but is secreted from LNCaP cells with an inverted topology. We demonstrate that Cavin1 expression is a critical regulator of Cav1 release, demonstrate intracellular Ca^2+^ levels affect secretion, and isolate and characterise the novel secreted vesicles from LNCaP cells; here termed C-exosomes (Caveolin-exosomes). We show by EM that C-exosomes are regular spherical structures of ∼40 nm in diameter that are highly enriched in Cav1 as demonstrated by tandem mass spectrometry proteomics analysis. Using single molecule fluorescence spectroscopy, we reveal that each C-exosome released from LNCaP cells consists of 50-60 caveolin molecules. Finally, we provide direct evidence that the release of C-exosomes from LNCaP cells stems from a non-classical autophagy-based mechanism. These observations identify a non-conventional pathway for the release of a novel antibody-accessible Cav1 vesicle which have important implications for prostate cancer.

## RESULTS

### Cav1 is secreted in an antibody-accessible form from LNCaP cells

Antibody-mediated neutralization of Cav1 has been demonstrated to inhibit PCa disease progression^12, 17^. This is inconsistent with a conventional exosomal release as Cav1 epitopes secreted within exosomes should be concealed within the lumen of the vesicle and inaccessible for antibody neutralization. To date Cav1 secretion has been most extensively studied in PC3 cells where Cav1 is known to co-fractionate with protein markers of conventional exosomes^14, 25^. Intriguingly, LNCaP cells expressing Cav1 have been shown to recapitulate the action of a bioactive form of Cav1 implicated in prostate cancer progression^19^ but it is currently unknown how Cav1 secretion occurs and what topology Cav1 adopts when released from these cells. To interrogate the pathway of release we first analysed the secretion of Cav1 from both PCa cell lines.

LNCaPs have variable levels of endogenous Cav1 expression; high passage LNCaP cells have been shown to have elevated expression and secretion of Cav1, yet low passage LNCaPs in the same study were shown to have no endogenous Cav1 expression^12^. Therefore, to evaluate the pathway of Cav1 secretion from LNCaP cells we have made use of a transient overexpression system to reliably study caveolin secretion as has been used in other studies *in vitro* and *in vivo*^11, 19^. As PC3 cells express and secrete endogenous Cav1 at reliably high levels we have performed the subsequent comparisons between untransfected PC3s and LNCaP cells transiently expressing Cav1.

Secreted vesicles were isolated from conditioned media by sequential centrifugations at 1,500 rpm for 5 min, 4,500 rpm for 20 min, and 14,000 x *g* for 35 min to remove cellular debris, and then ultracentrifugation at 100,000 x *g* for 60 min. To confirm that this purification method resulted in a sufficiently clean preparation of exosomes, we first analysed secreted vesicles from PC3 cell conditioned media by western blot analysis. The pelleted fraction (P100) from PC3 cells demonstrated abundant Cav1 protein (Fig. 1A) without contamination from other cellular compartments. Western blotting confirmed that protein markers of the Golgi complex, the nucleus, the cytoplasm and endosomes were absent (Fig. 1A), however, β-actin, α-tubulin, Flot1 and Cav1 (Fig. 1A) were all present as previously described for PC3 cell exosome perparations^14, 25, 31–33^. These data demonstrate this method is sufficient for generating a pure preparation of PCa cell exosomes.

**Figure 1.**
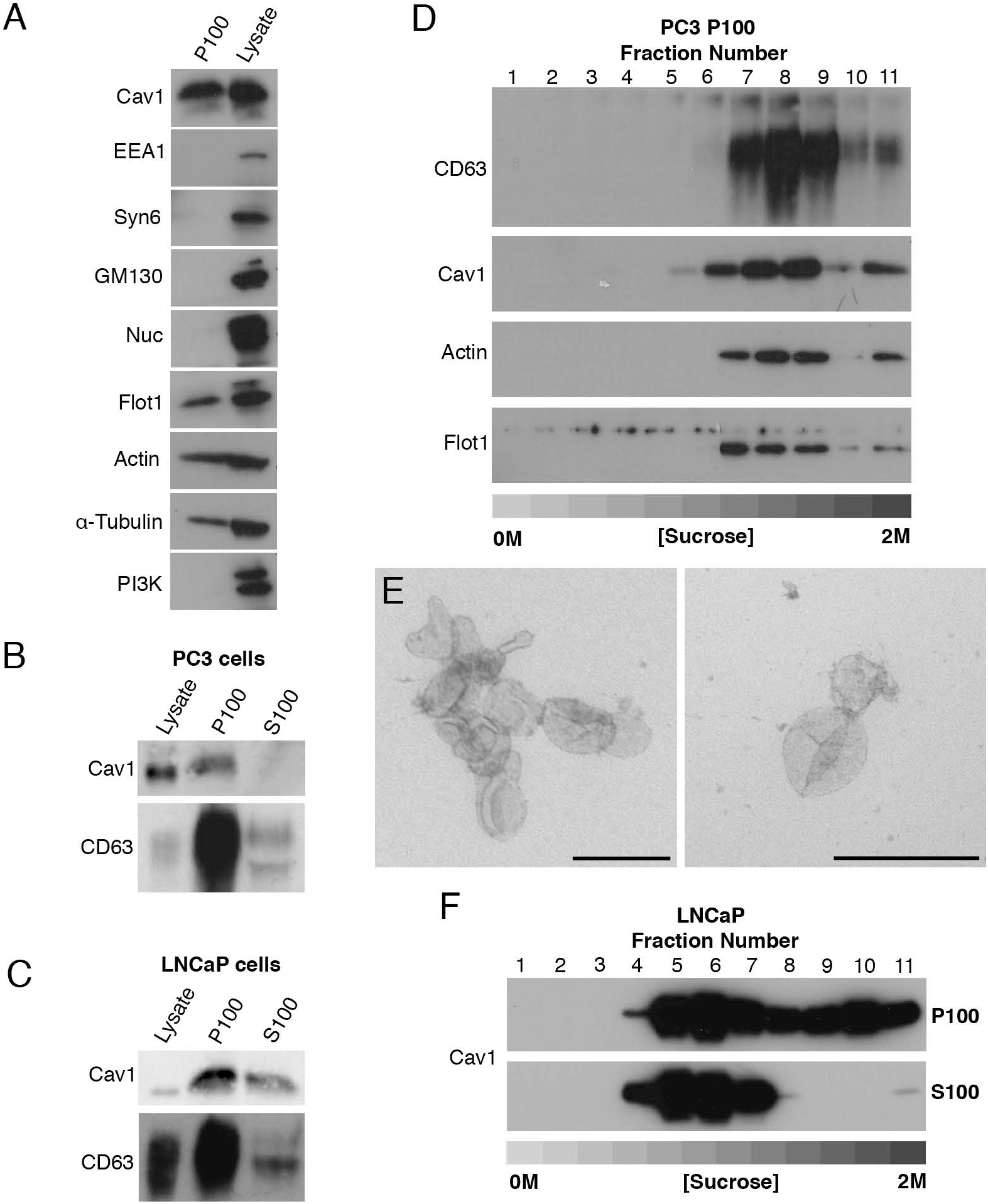
Cav1 is released in a novel form from LNCaP cells. A) Western blots of PC3 cells P100 fractions demonstrating the ultracentrigation purification method used in this study is not contaminated with subcellular protein markers. EEA1, Syntaxin-6, GM130, Nucleoporin and Phosphoinositide 3-kinase were absent from the P100 fraction in PC3 cells. Protein markers of exosomes including Flotillin-1, β-Actin and α-tubulin were detected by western blot. B) Western blot analysis and quantification of released Cav1 in P100 and S100 fractions. CD63 and Cav1 were enriched in the P100 fraction whereas Cav1 was absent from the S100 fraction. C) Western blot and quantification of the release of Cav1 from LNCaP cells transiently expressing Cav1. A proportion of Cav1 is observed in both the P100 and S100 fractions after ultracentifugation. D) Western blot of sucrose gradients from 0M to 2M of PC3 P100 fractions showing Cav1 fractionates at higher densities and co-fractionates with exosomal protein markers. E) EM of sucrose gradient fractions stained with 0.4% uranyl acetate mounted in 2% methyl cellulose. Peak exosomal morphology correlated with peak Cav1 protein levels. Left panel: fraction 7, right panel: fraction 8. Scale bars; 200 nm. F) Sucrose gradients of P100 and S100 fractions isolated from LNCaP cells demonstrating Cav1 is highly abundant in the S100 fraction consistently and fractionates at lower densities compared to Cav1 present in the P100 fraction and Cav1 secreted from PC3 cells. All western blots are representative blots chosen from three independent replicates.

As previous studies have demonstrated differences between the biophysical properties of exosomes isolated from PC3 cells compared to LNCaP cells we next assayed the relative abundance of Cav1 secreted into the P100 and S100 fractions between these two cell lines. Cav1 and CD63 (a protein marker of multivesicular bodies and exosomes) levels were analysed by western blot. In PC3 cells the amount of Cav1 released was proportionally small in the P100 fraction compared to the total cellular level of expressed protein only representing 0.03% of total cellular levels; no Cav1 protein was detected in the S100 fraction (Fig. 1B). In contrast, LNCaP cells demonstrated a different profile for Cav1 secretion with a larger proportion of total Cav1 protein present in the released fractions; 1.55% of total cellular Cav1 was in the P100 fraction and 1.13% released into the S100 fraction (Fig. 1C). Given the difference in secretion profiles between released Cav1 from PC3 cells and LNCaP cells, we next dissected the biochemical properties of secreted Cav1 from each cell line in greater detail using sucrose gradient fractionation. Cav1 in the P100 fraction derived from PC3 cells demonstrated an abundance of Cav1 protein in higher density fractions with peak protein concentration (between steps 7 to 11). Moreover, Cav1 secreted from PC3 cells co-fractionated with protein markers of exosomes and peak Cav1 levels closely correlated with peak exosomal morphology by electron microscopy (EM) (Fig. 1D and E). In contrast, sucrose gradient centrifugation demonstrated a significantly different fractionation profile for Cav1 isolated from LNCaP cells. Cav1 demonstrated wide-ranging densities from (steps 4 to 11) in the P100 fraction with a constrained low-density peak observed between steps 4 to 7 in the S100 fraction (Figure 1F). A time course of Cav1 release from PC3 cells demonstrated the presence of serum did not impact upon the dynamics of Cav1 secretion into the media (Fig. S1) which indicates that PC3 cell secretion of Cav1 does not originate from Cav1-positive exosomes present in the serum. These data confirm that Cav1 present in the S100 fraction from conditioned media isolated from LNCaP cells is biochemically distinct from the exosomal form isolated from PC3 cells.

To determine if the Cav1 particle secreted from LNCaP cells resembles the bioactive form of Cav1 in prostate cancer, we analysed the topology of the protein using immunoprecipitation. Cav1 was successfully immunoisolated from the S100 fraction from LNCaP cells using a polyclonal antibody against the Cav1 N-terminus (Fig. 2A). The pulldown of Cav1 from the S100 fraction was not dependent on pre-treatment with Triton-X100 (TX100), indicating that Cav1 is present in the S100 fraction in an exposed form (Fig. 2A). To determine the topographic organisation of Cav1 released from these cells, we performed pulldown analyses with GFP-trap using an N-terminal YFP-Cav1 construct and a C-terminal Cav1-GFP construct. Both the N- and C-termini were available for pulldown by GFP-trap (Fig. 2B). To validate the orientation of Cav isolated from PC3 cells we immunoisolated exosomes from the media (after the 14,000 x *g* centrifugation step) using anti-CD63 and polyclonal anti-Cav1 antibodies. Cav1 was detected in immuno-precipitated CD63-positive exosomes (Fig. 2C). Strikingly, the co-immunoprecipitation of Cav1 with CD63 was dependent on an intact membrane, as detergent treatment resulted in the loss of Cav1 pulldown by the anti-CD63 antibody (Fig. 2C). We next performed immunoprecipitation of the P100 fraction using an antibody against the N-terminus of Cav1. We could not pull-down Cav1 from the P100 fraction in the absence of detergent treatment but, in the presence of β-octylglucoside (βoG) and TX100, Cav1 was successfully immunoisolated (Fig. 2D). This strongly argues Cav1 is not available on the external leaflet of the exosome for antibody binding when secreted from PC3 cells but is secreted in an antibody-accessible form in the S100 fraction from LNCaP cells. To further confirm these results we performed protease protection assays against the S100 fraction from LNCaP cells and the P100 fraction from PC3 cells. In PC3 cells, Cav1 was partially digested only when detergent-treated, (Fig. S2A) consistent with previous observations^34^. In contrast, the recognition epitope of Cav1 was completely absent after incubation with ProK in the absence of detergent treatment in LNCaP S100 preparations (Fig. S2B). The complete loss of the recognition epitope after ProK treatment suggests the N-terminus of Cav1 may adopt an as yet undescribed topology in the membrane.

**Figure 2.**
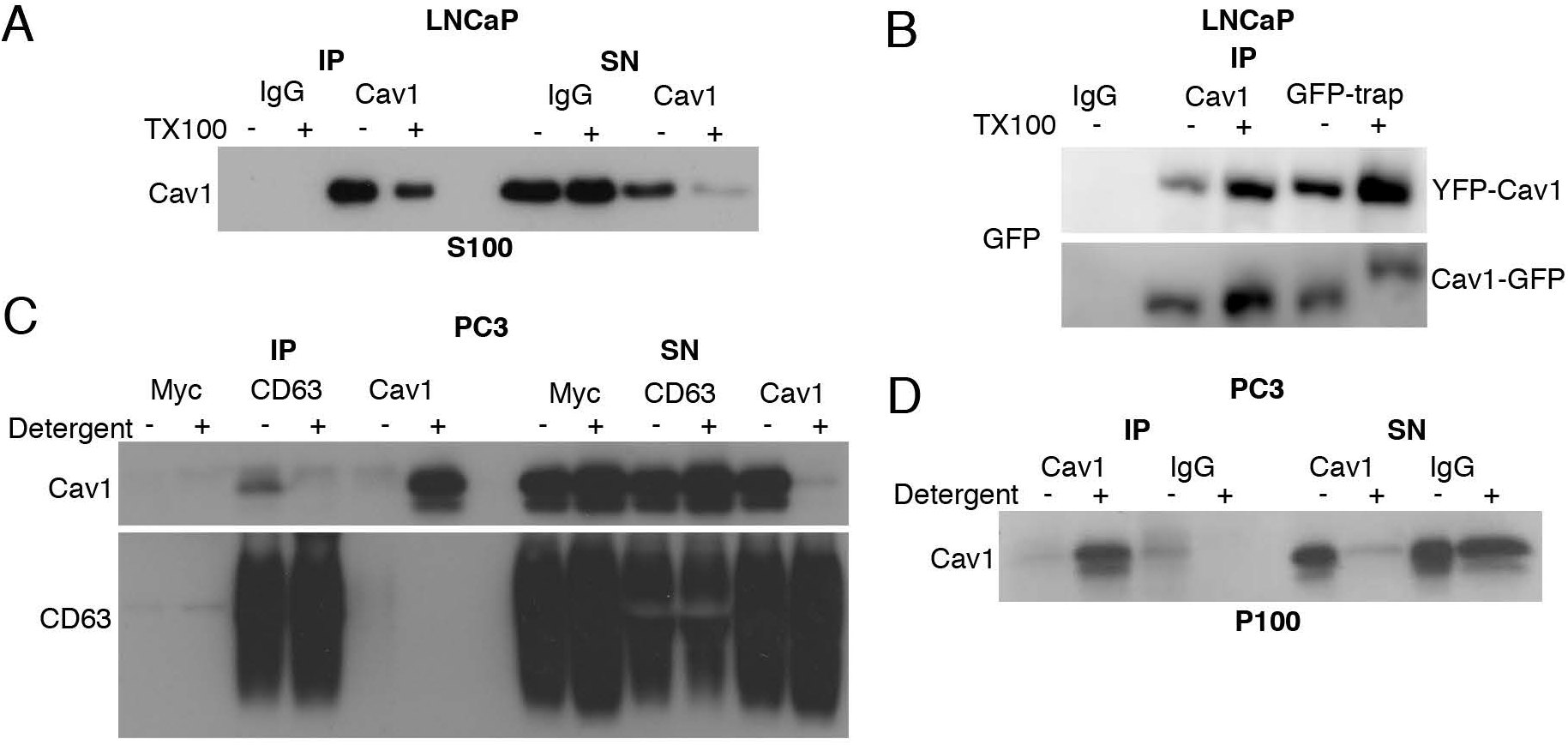
LNCaP cells release Cav1 in an antibody-accessible form. A) Western blot of an immunoprecipitation of Cav1 from the LNCaP S100 fraction with an α-Cav1 antibody demonstrating Cav1 can be immunoisolated in the absence of detergent. B) Cav1 can be pulled down by GFP-trap binding to fusion tags at both the N- and C-termini of Cav1 without detergent treatment. C) Immunoprecipitation of PC3 cell culture medium with an α-CD63 antibody results in the pulldown of Cav1 in the absence of detergent whereas immunoprecipitation with an α-Cav1 antibody does not result in the isolation of Cav1 unless pre-treated with a detergent. D) Western blot demonstrating pulldown of Cav1 from the PC3 P100 fraction is dependent on detergent treatment. All western blots are representative blots chosen from three independent replicates.

Taken together these data suggest that Cav1 is released from LNCaP cells in an atypical form, which we now term C-exosomes (Caveolin1-exosomes), that biophysically and topologically resemble the pro-angiogenic particles proposed to have autocrine and paracrine functions in prostate cancer models^12, 19–21, 35^.

### Cavin1 expression and caveolae formation inhibit the secretion of Cav1 from PCa cells

The conventional release of exosomes from the cell involves the sorting of proteins from the endosome into ILVs, the formation, fission and accumulation of ILVs within MVBs, and the subsequent fusion of MVBs with the PM. To gain a more detailed understanding of the pathway underpinning Cav1 release in LNCaP cells, we analyzed the role of Ca^2+^-mediated membrane fusion. The calcium ionophore, ionomycin, was utilised to determine how intracellular Ca^2+^ levels affected Cav1 secretion^36^. LNCaP cells secreted more Cav1 into the S100 fraction when treated with ionomycin in a time-dependent manner (Fig. 3A and B). Increased Cav1 levels were observed after only 10 minutes of treatment. A similar dependence on Ca^2+^-mediated membrane fusion was also observed in PC3 cells (Fig. S3A and B).

**Figure 3.**
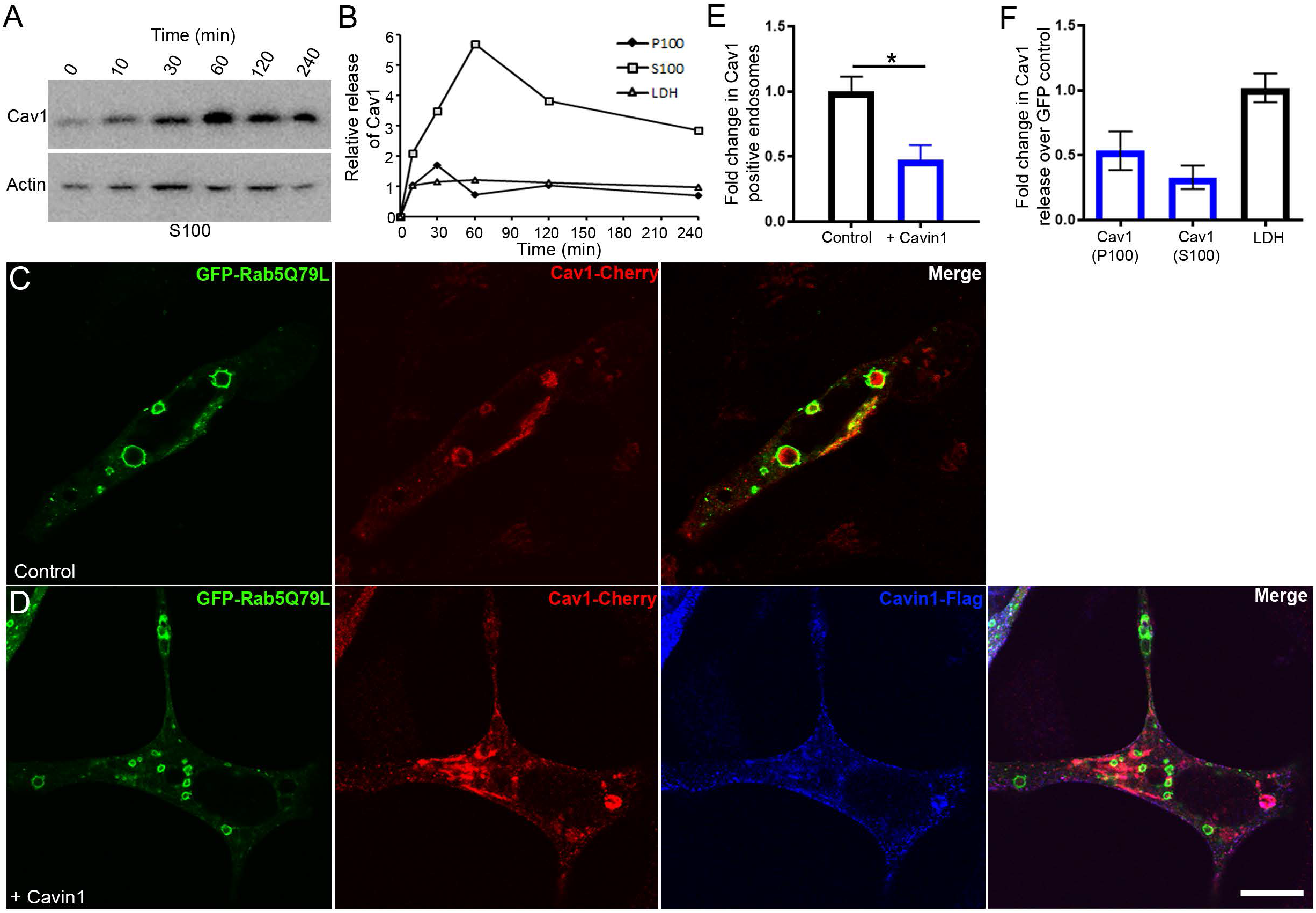
Regulated release of Cav1 from LNCaP cells. A) Western blot of Cav1 release from LNCaP cells stimulated with ionomycin. Cells were treated with 500 nM ionomycin in serum free media for various times (0 to 4 h). B) Peak release of Cav1 was observed after 1 h of treatment. Preferential release of Cav1 in the S100 fraction (compared to the P100 fraction) was observed and did not correlate with the release of LDH. A similar stimulation was observed for PC3 cells (see Fig. 3 Supplement - Figure 1A and B). C) LNCaP cells expressing GFP-Rab5Q79L; Cav1 is sorted into Rab5-positive endosomes in the absence of Cavin1 expression. D) Cavin1 expression sequesters and stabilises Cav1 at the PM within caveolae. E) Quantification of Cav1-positive GFP-Rab5Q79L endosomes demonstrates the expression of Cavin1 significantly reduced the internal pool of Cav1. Statistical significance was determined by two-tailed t tests; p = 0.0011, n = 3, error bars represent SEM. F) Quantification of western blots of the P100 and S100 fractions of Cavin1 expressing LNCaP cells compared to GFP expression alone. Cavin1 resulted in a reduction in Cav1 release in both fractions without altering LDH levels. n = 4; error bars represent SEM. We also confirmed that Cavin1 expression in PC3 cells (also devoid of Cavin1 expression^6^) reduced internalisation and secretion of Cav1 (see Fig. S3).

In view of the antibody-accessible topology of Cav1 released from LNCaP cells, we next interrogated the mechanistic requirements underlying the release of this novel form. To dissect the requirements for the release of Cav1 we analysed the importance of Cavin1 expression as Cavin1 is structurally required for caveolar biogenesis/stability^6^, regulates Cav1 internalisation^9^ and is not expressed in LNCaP or PC3 cells^11, 16^. LNCaP cells were transiently transfected with the GFP-Rab5Q79L point mutant, which is a GTPase-deficient Rab5 that stimulates early endosome fusion^37^. This point mutant results in the formation of larger endosomes that are readily resolvable by confocal fluorescence microscopy. Using this assay, we performed a quantitative assessment of the relative internalized pool of Cav1 with and without Cavin1-Flag co-expression. The expression of Cavin1 significantly inhibited the internalisation of Cav1 in LNCaP cells (Fig. 3C, D and E) and PC3 cells (Fig S3C, D and E). These data suggest that expression of Cavin1-mediated sequestration of Cav1 at the PM inhibits intracellular accumulation.

We further tested if Cavin1 expression was sufficient to reduce the release of Cav1. We generated stable LNCaP cell lines expressing Cavin1-GFP or GFP alone with transient expression of Cav1-cherry. Cavin1-GFP expression (compared to GFP alone) resulted in a dramatic reduction in the release of Cav1-cherry in both the S100 (73% ± 9) and P100 (47% ± 15) fractions (n = 4; quantified in Fig. 3F) suggesting that Cavin1 expression sequesters Cav1 within caveolae at the PM resulting in reduced secretion into the extracellular space. A similar reduction in Cav1 secretion was observed upon Cavin1 expression in PC3 cells by western blot analysis (Fig. S3F). Taken together these data indicate that intracellular Ca^2+^ levels affect caveolin release, and that Cav1 secretion is negatively regulated by Cavin1 expression.

### Molecular and ultrastructural analysis of Cav1 released from LNCaP cells

Our results have demonstrated an N-terminal YFP-tag is sufficient to immuno-isolate Cav1 from the S100 fraction of LNCaP cells. We used this observation to gain insights into the molecular composition and structure of C-exosomes to determine the origin of their release. Initial experiments successfully used GFP-trap beads to isolate YFP-Cav1 from S100 fraction of LNCaP cells, however, the yields were low. We went on to optimise purification with MBP-tagged GFP-trap, purified on amylose resin column, eluted with maltose buffer, and then concentrated in 100kD cut-off centricon filters. Proteomic analysis was performed on three biological replicates of YFP-Cav1 particles purified from LNCaP conditioned media, compared to YFP expression alone. A total of 453 proteins were identified across the 6 samples (Table S1), most not consistently detected across replicates, indicative of non-specific interactions. Strikingly, only 2 proteins were significantly different (p<0.05) between YFP-Cav1 and YFP groups across the 3 replicates: Caveolin-1, as expected, and Synaptogyrin-2. This protein has previously been implicated in endocytic and synaptic vesicle formation^38, 39^. Importantly, the lack of commonly observed exosomal proteins such as Flotillin-1, Flotillin-2, or TSG101 further confirms that YFP-Cav1 particles are not released as conventional exosomes.

We further characterized the released particles by EM. YFP-Cav1 was isolated from conditioned media using method (ii) as this protocol yielded the largest amount of pure material. The particles (still with MBP tagged GFP-trap attached) were spherical in structure, uniform in diameter and morphology (Fig. 4A) measuring ∼36 nm from inner membrane to inner membrane (the outer diameter was not measured because it includes the MBP-GFP-trap bound to YFP-Cav1); smaller than conventional exosomes (∼100-200 nm) and plasma membrane caveolae (50 to 80 nm). To obtain a quantitative understanding of the number of Cav1 proteins per C-exosome we used fluorescence correlation spectroscopy (FCS)^40^. YFP-Cav1 positive particles were isolated and analysed under the same conditions as YFP alone (as a calibration factor). 100 curves of 10s were acquired for YFP and the diffusion time was plotted (Fig. 4B); the data show a narrow distribution of residence times centred at 95 μsec. YFP-Cav1 however demonstrated a broader distribution of residence times ∼800 μsec (Fig. 4C) approximately 7.5 times the size of the YFP monomer. Using these values, an approximation of particle size can be gained by comparison with the size of YFP (∼5.5 nm) at ∼40-45 nm, close to the measured diameter from negative staining EM.

**Figure 4.**
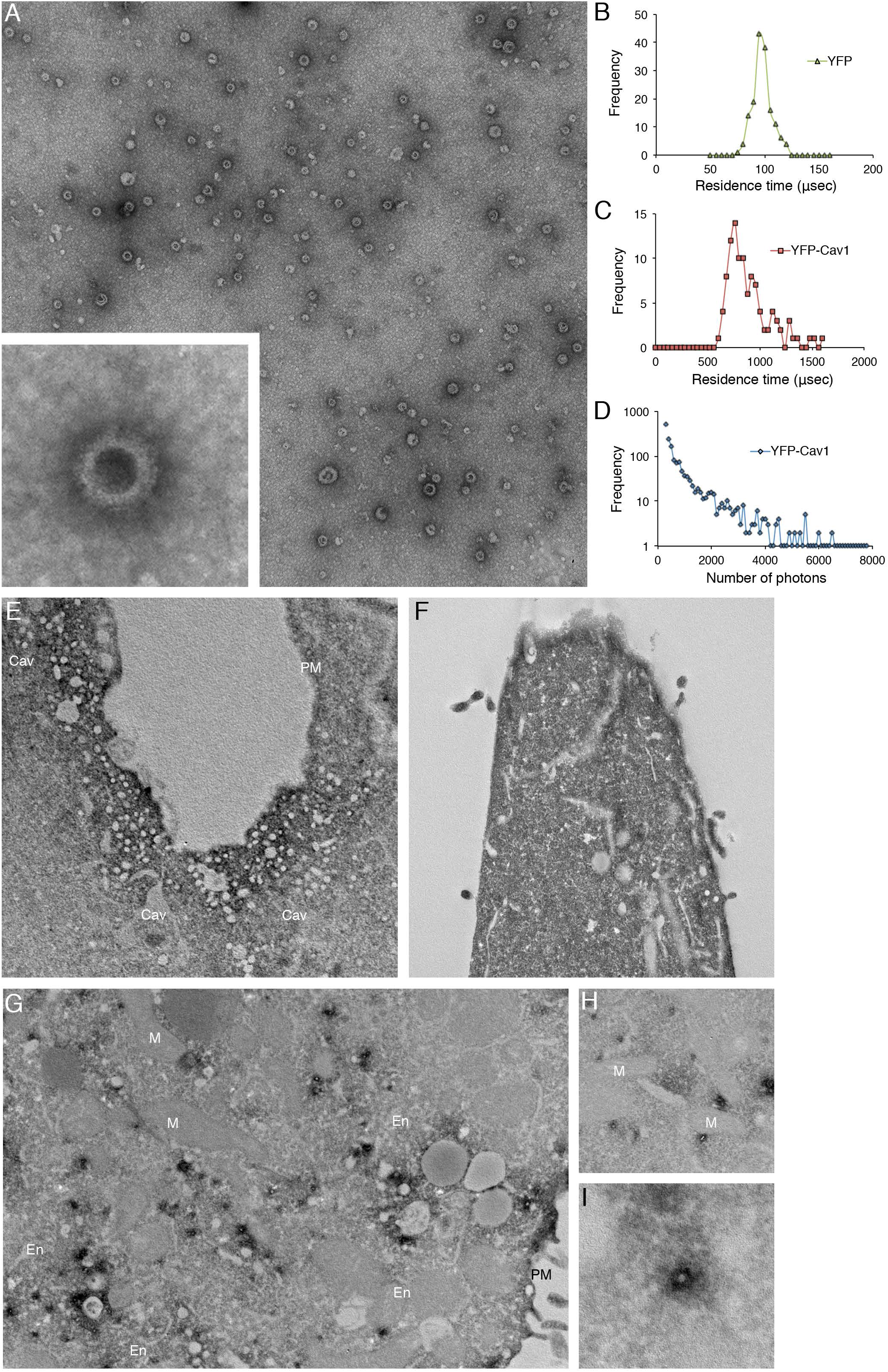
High-resolution analyses of C-exosomes and YFP-Cav1 in LNCaP cells. A) EM of YFP-Cav1 isolated from LNCaP media demonstrates particles are regularly shaped vesicles. Inset shows high-magnification image of stained LNCaP particles with a prominent coat on the external leaflet, which includes the YFP fusion and GFP-trap with a maltose binding protein tag. Scale bar = 500 nm. B - D) Single molecule analysis of YFP-Cav1 released from LNCaP cells. Control analysis of YFP alone demonstrates a residence time of approximately 95 μsec. C) Analysis of the residence time of YFP-Cav1 demonstrates a 7.5-fold increase in the residence time compared to YFP alone. D) Plot of burst brightness against the number of bursts of that brightness demonstrates the maximum number of YFP-Cav1 molecules per vesicle to range between 50-60 proteins. E – I) EM analyses of YFP distribution in cells using APEX-GBP expressing YFP-tagged constructs. E) Co-expression of YFP-Cav1 and APEX-GBP demonstrates morphologically typical localisation of Cav1 in BHK cells at caveolae on the PM of expressing cells. Arrows denote the caveolae with enriched electron density by the association of APEX-GBP. Cav = caveolae, PM = plasma membrane, scale bar = 500 nm. F) The co-expression of YFP and APEX-GBP demonstrates a cytoplasmic distribution for YFP alone. Scale bar = 500 nm. G) Lower magnification image of an LNCaP cell transfected with APEX-GBP and YFP-Cav1. Arrows highlight small vesicles with very strong reaction products that were absent from BHK cells. Black Arrowheads denote Cav1 positive endosomes and red arrow demonstrates Cav1 localisation to the PM. H) YFP-Cav1 vesicles are regular in size. I) High-magnification image of an YFP-Cav1 shows vesicles are membrane bound. Scale bars = 500 nm. Electron micrographs are representative images; each LNCaP experiment was independently replicated three times.

Analysis of the predicted size distribution demonstrates the YFP-Cav1 particles released by LNCaP cells are highly defined in size and number when compared to random aggregation, which possesses broader distribution and diffusion values (> 10,000 μsec)^40^. We then used single molecule counting to estimate the number of proteins contained in the YFP-Cav1 particles^41^. Plotting the brightness of bursts against the number of bursts of that brightness, released YFP-Cav1 showed a large maximal amplitude of about 4,500 photons (Fig. 4D). As a single YFP protein can generate a maximum of 90 photons under the same conditions, this yields a relative value of approximately 50 to 60 units YFP-Cav1 proteins per C-exosome. These data demonstrate YFP-Cav1 is released from LNCaP cells as a regular, spherical vesicle comprising 50-60 proteins per particle in a form that is different from conventional exosomes and caveolae.

To further characterise Cav1 secreted from LNCaP cells, we performed EM with APEX-GBP; a modular method for the high-resolution detection of subcellular protein distributions^42^. APEX-GBP is an expression vector with a modified soybean ascorbate peroxidase tag^43^ linked to a high affinity GFP/YFP-binding peptide^44^. The APEX-tag generates an osmiophillic polymer when the diaminobenzoic acid (DAB) reaction is performed in the presence of H_2_O_2_; this insoluble polymer is contrasted by osmium tetroxide post-fixation which allows for the detection of any GFP- or YFP-tagged protein to an approximate 10 nm resolution^42^. To confirm the expression of YFP-Cav1 and APEX-GBP is non-disrupting, we first expressed these constructs in the non-PCa line, baby hamster kidney (BHK) cells, which endogenously express Cavin1. The co-transfection of YFP-Cav1 with APEX-GBP resulted in a normal Cav1 distribution with enriched electron density at the PM, specifically at caveolae with minimal electron density at intracellular structures (Fig. 4E). We next examined the expression of YFP alone or YFP-Cav1 with APEX-GBP in LNCaP cells. YFP co-transfection with APEX-GBP resulted in soluble/cytoplasmic electron density with no enrichment at membrane compartments (Fig. 4F). YFP-Cav1 co-expression with APEX-GBP in LNCaP cells resulted a broad distribution of subcellular localisations with Cav1 occasionally detected at flat PM (red arrow) and at endosomes (black arrowheads). Intriguingly, YFP-Cav1 was highly abundant at small (∼31±8 nm internal diameter) intracellular vesicular structures of regular size and shape that were completely disconnected from the PM and other membrane-bound cellular compartments. The electron density generated by the APEX-tag and the DAB reaction demonstrated that the topology of YFP-Cav1 in these small particles was consistent with exposure on the cytoplasmic face of these vesicles (Fig. 4G-I) and were morphologically similar in size and shape to those imaged by negative staining.

The distribution of YFP-Cav1 was markedly different in PC3 cells. Cav1 was abundant on flat PM but predominantly decorated the cytoplasmic face of endosomal compartments (Fig. 5B: white arrows) and with striking density inside intraluminal vesicles accumulating within endosomes (Fig. 5B, C and E: white arrowheads). Furthermore, profiles consistent with endosomes fusing with the PM to release YFP-Cav1:APEX-GBP-rich vesicles were regularly observed (Fig. 5E) as well as exosomes positive for YFP-Cav1 on the external surface of PC3 cells (Fig. 5D and E).

**Figure 5.**
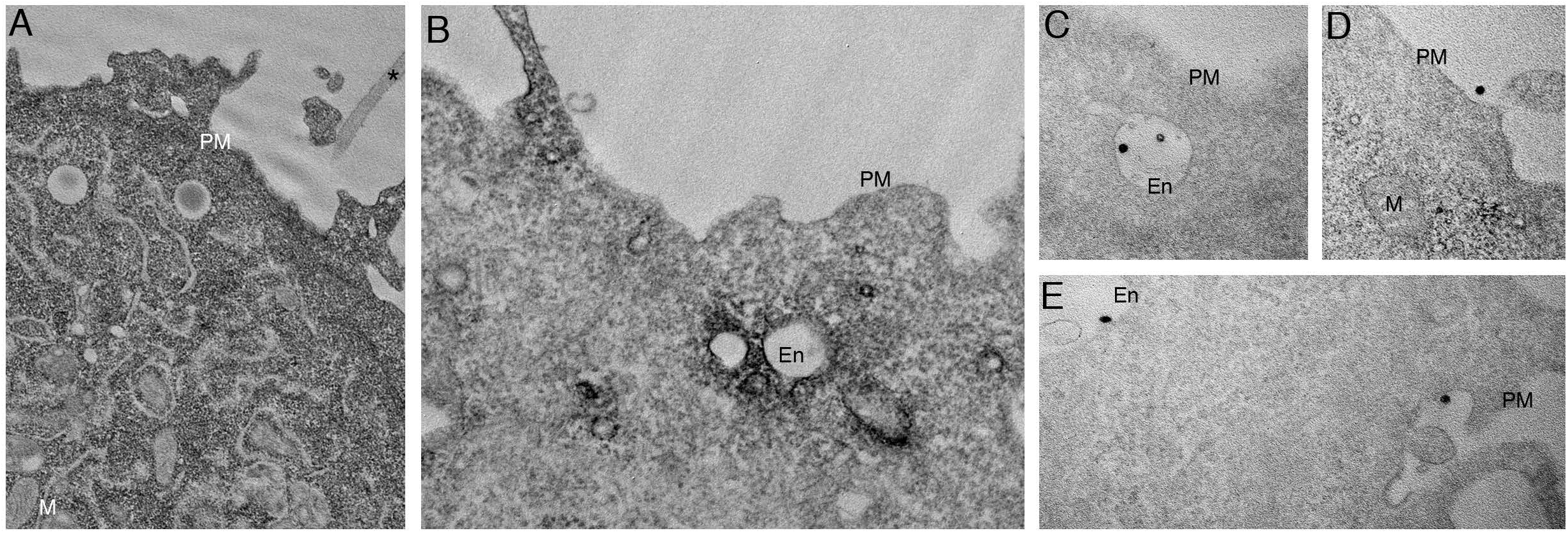
Cav1 expression in PC3 results in predominantly endosomal localisation by APEX-GBP and EM. EM analysis of YFP distribution in cells using APEX-GBP expressing YFP-tagged constructs. A) The co-expression of YFP and APEX-GBP results in soluble reaction product localised to the cytoplasm of transfected cells. Asterisk denotes the electron density of an untransfected adjacent cell, PM = plasma membrane, M = mitochondria, scale bar = 500 nm. B) Electron micrograph of a PC3 cell co-transfected with YFP-Cav1 and APEX-GBP. Electron density shows significant enrichment of Cav1 at endosomes (white arrows), the plasma membrane (black arrow) and less frequently small vesicular structures in the cytoplasm (black arrowhead). En = Endosome, PM = plasma membrane, scale bar = 500 nm. C – E) ILVs (white arrowheads) and secreted ILVs/exosomes (red arrowheads) highly enriched with YFP-Cav1 and APEX-GBP inside endosomes and outside the cell. Scale bars = 500 nm. Electron micrographs are representative images; each PC3 experiment was independently replicated two times.

In view of these observations we speculated that Cav1 may drive the formation of 30-40 nm C-exosomes in the LNCaP cytoplasm through the protein’s ability to sculpt membranes^45^. We interrogated this model by comparing secretion of WT Cav1 with secretion of the Cav1S80E point mutant, which inhibits Cav1 membrane sculpting by reducing membrane affinity in a bacterial system^34, 45^ and cholesterol binding in a mammalian system^46^. The expression of the S80A point mutant, which increases cholesterol affinity^46^ and is non-disrupting in the bacterial system^45^, resulted in a modest increase in the release of Cav1S80A compared to the release of WT Cav1 (Fig. 6A and B). However, the expression of the Cav1S80E point mutant resulted in a two-fold reduction in released Cav1 (Fig. 6A and C). Confocal microscopy demonstrated that the S80E mutant was exclusively localised to the Golgi complex (as determined by co-localisation with GM130), whereas the WT construct was only partially co-localised with this organelle marker (Fig. 6D and E). WT Cav1 was efficiently sorted into endosomes, as determined by co-transfection with GFP-Rab5Q79L, but Cav1S80E was unable to accumulate within Rab5Q79L-positive compartments. Additionally, the expression of Cavin1 did not result in the redistribution of Cav1S80E to the cell surface unlike the WT Cav1 (Fig. 6F-I). These data suggest that cholesterol binding is essential for the trafficking of Cav1 out of the Golgi complex for secretion into the extracellular space. To confirm this, we utilised APEX-GBP to examine the distribution of Cav1S80E at the EM level. The S80A mutant efficiently generated the cytoplasmically-localised ∼35 nm vesicles (Fig. 6J), was lowly abundant at cell surface and on endosomes similar to the WT Cav1 (Fig. 4G-I). Cav1S80E however, was unable to efficiently generate these vesicles and remained almost completely associated with the Golgi complex (Fig. 6K).

**Figure 6.**
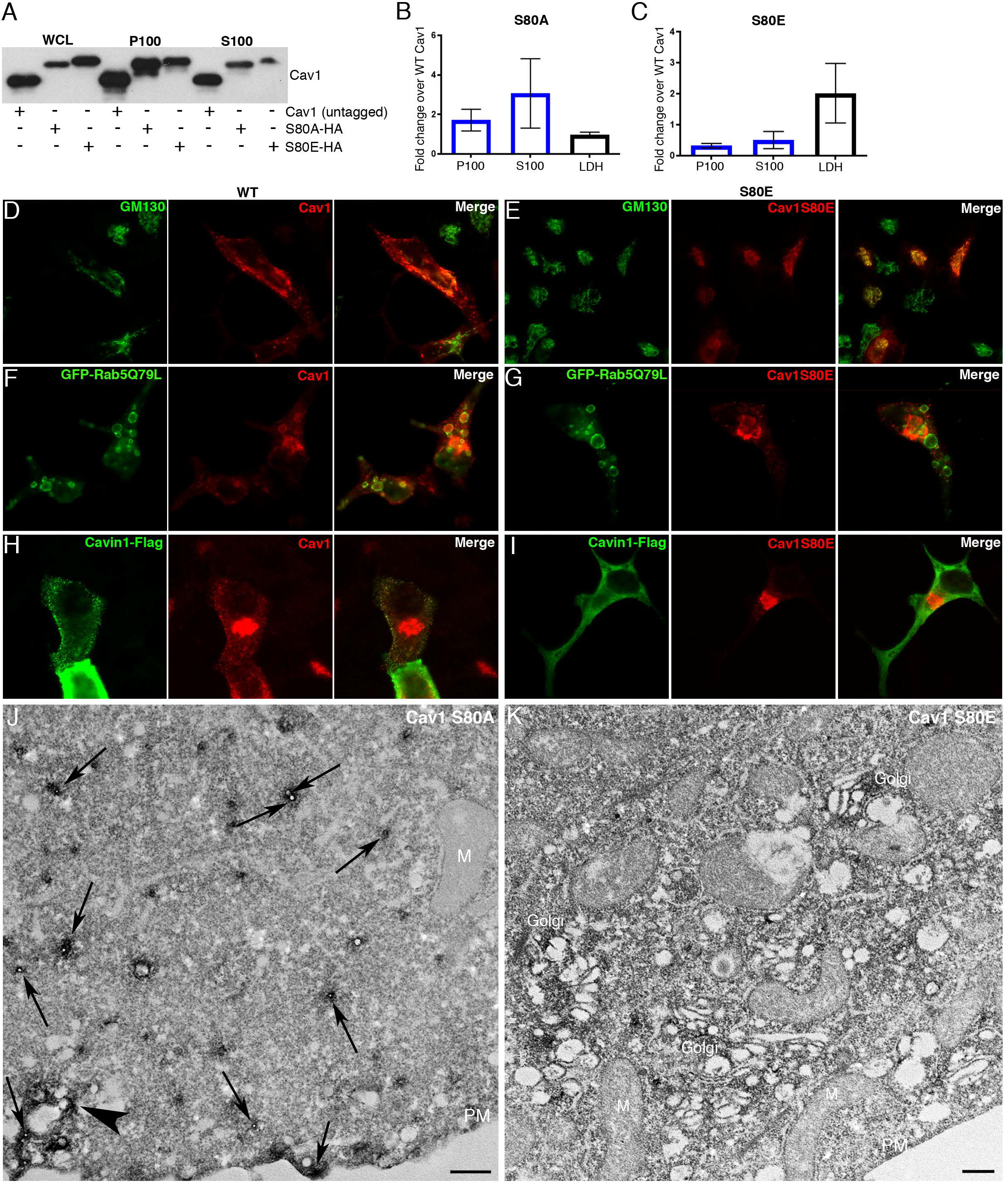
Cav1 mutants are differentially released from LNCaP cells. A) Western blot of the S100 and P100 fractions of Cav1 released from S80A and S80E expressing LNCaP cells. B) A relative increase in the release of the S80A in both the P100 and S100 fractions was observed compared to WT (n = 3). C) A relative reduction in the release of the S80E mutant was observed compared to WT. While LDH levels were increased with the expression of the Cav1S80E mutant, this would likely result in a corresponding increase in the non-specific release of. Despite this, a two-fold reduction in secreted Cav1 levels were observed. D) WT Cav1 partially co-localises with GM130 at the Golgi complex. E) The S80E mutant is almost exclusively localised to the Golgi complex. F) WT Cav1 is sorted into GFP-Rab5Q79L positive compartments. G) S80E mutant is not efficiently sorted into GFP-Rab5Q79L-positive endosomes H) Cavin1-Flag expression stabilises Cav1 in punctate structures at the PM of LNCaP cells. I) Cavin1-Flag expression does not stabilise Cav1 at the surface of LNCaP cells and Cavin1 remains cytoplasmic/soluble when co-transfected with the S80E mutant. J) LNCaP cells demonstrating the S80A point mutant efficiently generated C-exosome precursors in the cytoplasm. C-exosomes = black arrows. Scale bar = 500 nm. K) LNCaP cells expressing YFP-Cav1S80E mutant construct. The point mutation predominantly is localised to the Golgi complex and is inefficient in generating small Cav1-rich vesicles. Scale bar = 500 nm. Fluorescent images are representative images from three independent replicates. Electron micrographs are representative images; each LNCaP experiment was independently replicated three times.

Taken together, the characterisation of C-exosomes released by LNCaP cells indicate the particles isolated from the S100 fraction (i) are not caveolae released from lysed cells, (ii) are not conventional exosomes equivalent to Cav1 released by PC3 cells, (iii) have an inverted topology consistent with the bioactive form of Cav1 involved in prostate cancer, and (iv) mutations in Cav1 that decrease membrane sculpting cause retention in the Golgi complex and inhibit Cav1 secretion.

### Autophagy is critical for the release of Cav1 from LNCaP cells

Autophagy has been previously implicated in the release of proteins through atypical pathways^47–51^ and several studies have linked degradation of Cav1 to autophagy^52–55^. Our EM studies demonstrated abundant small vesicles positive for Cav1 in the cytoplasm of expressing cells. We hypothesised these vesicles may be secreted after engulfment by the maturing autophagosome and then released from the cell^47–51^. Therefore, we first performed immunofluorescence and labelled endogenous LC3B, a marker of autophagosomes^56^, in YFP-Cav1 expressing LNCaP cells. YFP-Cav1 showed co-localisation with LC3B-positive puncta (Fig. 7A), suggesting the sorting of Cav1 into the autophagosomes in LNCaP cells.

**Figure 7.**
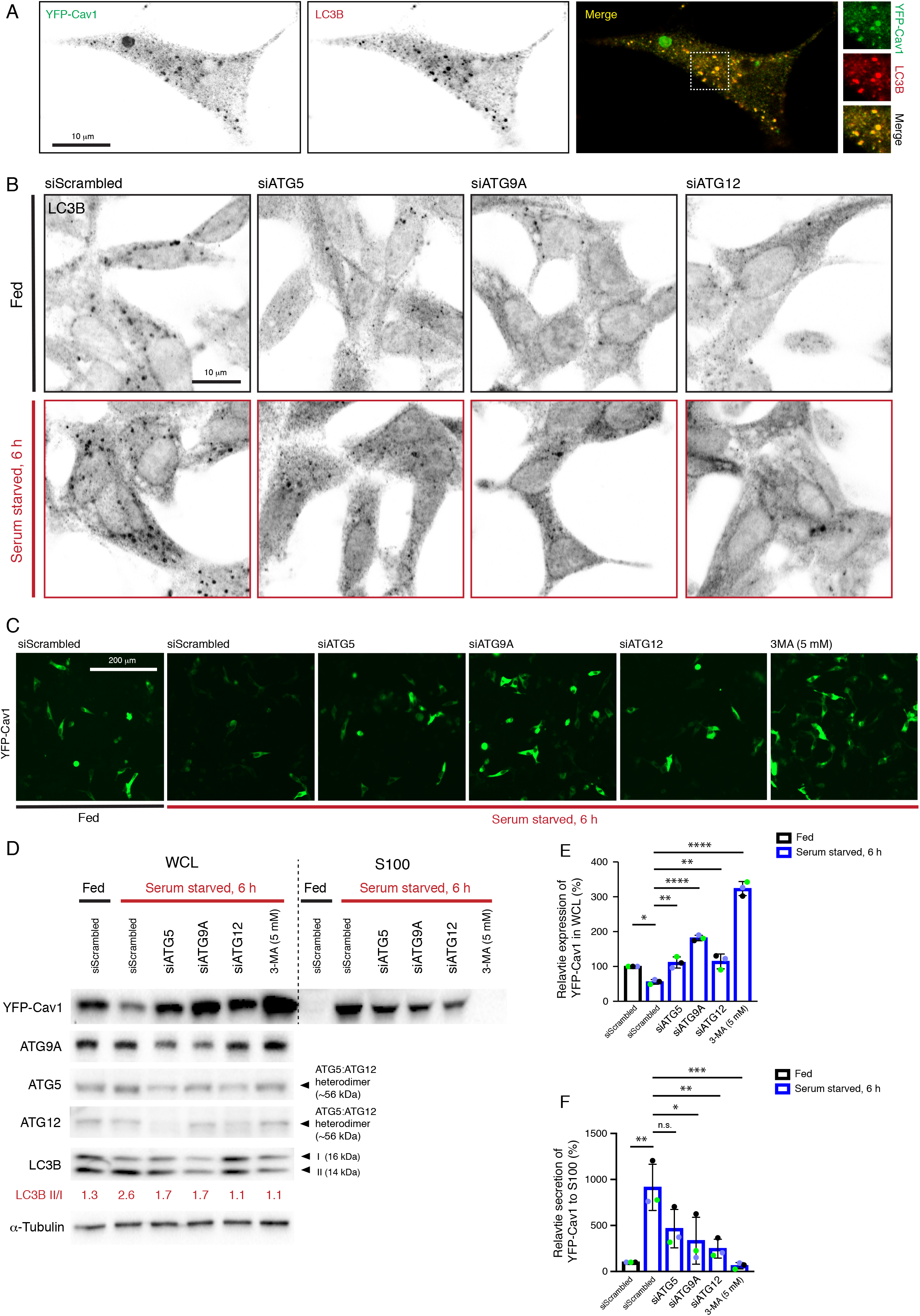
Cav1 secretion from LNCaP cells is mediated by an autophagy-dependent pathway. A) Representative confocal microscopy images from three independent experiments showing co-localisation between YFP-Cav1 and LC3B in LNCaP cells. Endogenous LC3B was immuno-stained with rabbit anti-LC3B primary antibodies followed by a secondary Alexa555 fluorescent labelling. Single channel images were converted to black and white and the contrast was inverted. The enlarged region demonstrates the overlapping distribution between YFP-Cav1 and LC3B. Scale bar, 10 μm. B) LNCaP cells were serum starved for 6 h to induce autophagy. Autophagosome formation was indicated by the immunofluorescence for endogenous LC3B. Knockdowns of ATG5, −9A and −12 proteins inhibited autophagosome formation in both fed (upper panel) and serum starved (lower panel) LNCaP cells. Representative images of three independent experiments were converted to black and white and contrast inverted. Scale bar, 10 μm. C) Representative fluorescent images demonstrating a reduction of YFP-Cav1 expression levels in serum starved LNCaP cells. Knockdown of ATG5, −9A or −12 and 3MA (5mM, 6 h) treatment restored YFP-Cav1 expression levels in starved cells. Scale bar, 200 μm. D) Representative western blots showing YFP-Cav1 protein levels in whole cell lysates (WCL) and S100 fractions of LNCaP cells equivalently transfected with YFP-Cav1 (10 μg DNA per 150 × 25 mm dish). The knockdown efficiency was assessed by the detection of ATG5, −9A and −12 protein levels. The ratio of LC3B II to LC3B I (LC3B II/I) was calculated as an indicator of autophagic levels in each group. An increase in the LC3B II/I ratio of the serum-starved siScrambled cells (2.8) compared to the fed siScrambled cells (1.3) indicates the induction of autophagy. E) Densitometry analysis of Western blots shown in (D) demonstrate that the co-transfection with ATG5, −9A or −12 siRNAs and 3MA treatment significantly (siATG5 starved vs. siScrambled starved: p = 0.0047; siATG9A starved vs. siScrambled starved: p < 0.0001; siATG12 starved vs. siScrambled starved: p = 0.0031; 3MA starved vs. siScrambled starved: p < 0.0001) rescued the downregulation of YFP-Cav1 (siScrambled fed vs. siScrambled starved: p = 0.0246) in the WCL of serum-starved cells compared to the fed cells. One-way ANOVA was performed for statistical analysis. F) The secretion (%) of YFP-Cav1 into the S100 fraction from LNCaP cells was significantly upregulated (siScrambled fed vs. siScrambled starved: p = 0.0011) upon serum starvation. The blockage of serum starvation-induced autophagy via ATG5, −9A, −12 knockdowns or 3MA treatment downregulated the secretion levels of YFP-Cav1. Statistically-significant effects (one-way ANOVA) were observed in ATG9A siRNA (siATG9A starved vs. siScrambled starved: p = 0.0152), ATG12 siRNA (siATG12 starved vs. siScrambled starved: p = 0.0056) and 3MA (3MA starved vs. siScrambled starved: p = 0.0007) treated groups.

Next, we utilised small interfering RNA directed against autophagy related (ATG) proteins ATG5, ATG9A and ATG12 that function at different stages of autophagy to investigate a potential role in Cav1 clearance. ATG9A is an upstream autophagic factor that is responsible for the delivery of the lipids and proteins required for autophagosome formation^57, 58^, while ATG5 and ATG12 play critical roles in a later stage of autophagy through forming a heterodimer that promotes autophagosome elongation^59^ and binds directly to Cav1^53^. As indicated by LC3B puncta (Fig. 7B), serum starvation for 6 hours effectively induced autophagosome accumulation. Knockdown of ATG5, ATG9A or ATG12 inhibited autophagosome formation under both nutrient (Fig 7B; upper panel) and serum starved (Fig 7B; lower panel) conditions. The modification of the LC3B cytoplasmic form (LC3B-I) to its membrane-bound form (LC3B-II) is essential for autophagosome maturation^56^. Therefore, LC3B lipidation was further detected to assess the autophagic levels. A two-fold increase in LC3B II/I ratio was observed in controls after a 6-hour serum starvation (Fig. 7D, n=3). Cells transfected with siRNAs against selected ATG proteins exhibited decreased LC3B-II/I ratios (Fig.7D), suggesting reduced autophagic levels upon serum starvation in those cells. The interruption of autophagy by knockdown of ATG proteins resulted in a dramatic increase in cellular YFP fluorescence (Fig. 7C) and significant upregulation of YFP-Cav1 protein levels in the whole cell lysates (WCL) from serum starved LNCaP cells (Fig. 7D and E). In addition, the treatment of an autophagy inhibitor, 3-methyladenine (3MA), was included as a positive control and exhibited the most significant effect on rescuing cellular YFP-Cav1 expression levels (3.21 fold) following serum starvation (Fig. 7C-E). These data demonstrate that autophagy is essential for the clearance Cav1 from LNCaP cells.

Next, we investigated the effect of autophagy on the release of Cav1 into the extracellular space. YFP-Cav1 expressing LNCaP cells transfected with siRNAs targeting ATG proteins or treated with 5 mM 3MA were incubated in serum free media for 6 hours to stimulate autophagy and the S100 fractions were then isolated. Western blot assays together with densitometry analysis revealed a significant upregulation (9.12 fold) of YFP-Cav1 secretion into the S100 fraction from control cells cultured in serum free medium for 6 hours (Fig. 7D and F). Autophagy inhibition by knockdowns of selected ATG proteins or 3MA treatment caused marked reduction in YFP-Cav1 secretion from starved LNCaP cells (Fig. 7D). Quantification showed significant downregulation in YFP-Cav1 levels in the S100 fractions of media isolated from ATG9A (−0.64 fold) or ATG12 (−0.73 fold) depleted cells, as well as 3MA treated cells (−0.93 fold), compared to controls (Fig. 7F). Despite no significance, knockdown of ATG5 led to a decrease of 49% of YFP-Cav1 secretion into the S100 fraction compared to controls (Fig. 7F).

Taken together, the results argue that autophagy is the major pathway for Cav1 secretion in LNCaP cells. We propose that Cav1-induced vesicles are engulfed by autophagosomes and some of this content is released into the extracellular space as C-exosomes (Schematically depicted in Fig. 8).

**Figure 8.**
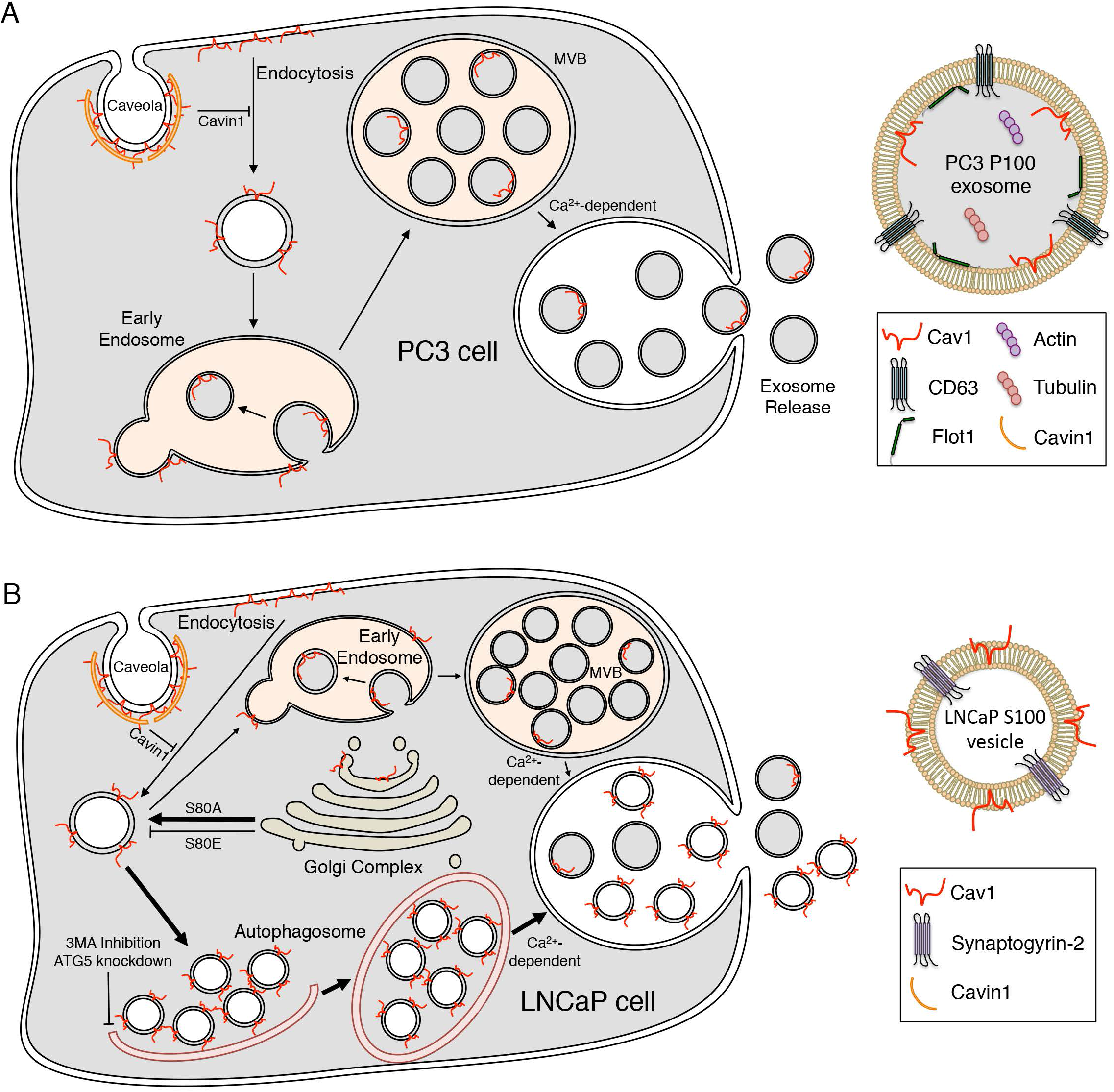
Cav1 secretion from PCa cells. A) Schematic summary of the exosomal release of Cav1 from PC3 cells. B) Schematic summary of the autophagy-based secretion of Cav1 from LNCaP cells.

## DISCUSSION

Serum Cav1 detection in PCa correlates with cancer stage^60^, cancer grade^61^, angiogenesis^18^, and poor patient outcomes^21, 62^. However, the mechanism that underlies this secretion has, to date, been poorly understood. Utilizing a variety of biochemical, EM and fluorescence microscopy-based techniques we have characterised the release of Cav1 from LNCaP cells and determined that Cav1 is released in a novel form through an autophagy-based pathway.

Advanced PCa possess an unconventional expression profile with high levels of Cav1 in the absence of Cavin1 expression^6, 11^. This is particularly interesting as loss of Cavin1 protein in cells and animals, and loss of function mutations in Cavin1 in human patients, consistently results in a significant reduction in caveolin protein levels^6, 7, 63, 64^. These observations suggest that understanding how PCa cells retain Cav1 expression for secretion into the extracellular space is critical for understanding Cav1’s role in PCa. Clearly, Cav1 can be released within exosomes through a conventional pathway; this is the predominant pathway for Cav1 release from PC3 cells. However, we now show that Cav1 is released from LNCaP cells with the opposite topology (summarized in Fig. 8). This is backed up by numerous observations from the literature that show bioactive Cav1-positive vesicles are released in an LNCaP-like topology with a biochemical profile that is distinct from conventional exosomes; Cav1 particles can be neutralised by anti-Cav1 antibodies which in turn can inhibit disease progression in models of PCa^12, 17^. These observations closely align with our characterization of the topology Cav1 released by LNCaP cells and suggest the orientation of Cav1 in the membrane may be a critical factor in disease progression and bioactivity.

Cav1 is oriented in the cell such that the N- and C-termini face the cytoplasm with no portion of the protein exposed to the extracellular milieu^65^. Our studies have confirmed that despite opposing secreted topologies, both PC3 cells and LNCaP cells possess Cav1 that exclusively faces the cytoplasm. How then can Cav1 be released from LNCaP cells with both cytoplasmic termini exposed? One possibility is that LNCaP particles are a consequence of lysed cells releasing caveolae into the media; our observations however argue against this possibility. First, LNCaP cells possess little to no caveolae in the absence of Cavin1 expression (see reference ^11^ and Fig 4G). Second, the release of Cav1 appears to be a regulated pathway as ionomycin treatment stimulated secretion and 3MA treatment inhibited release and critically these treatments did not correlate with large changes to LDH levels in the media. Third, the biophysical density of Cav1 secreted from LNCaP cells is dissimilar to the density of caveolae but does closely correlate with Cav1 released in other non-caveola forms^19, 66^. Fourth, the number of caveolin proteins per C-exosome does not align with the number of caveolin proteins per caveolae^5^. Finally, the vesicles released from LNCaP cells in the S100 fraction are approximately half the diameter of caveolae isolated from the PM^67^. Our data argue that precursor C-exosomes can be engulfed and released into the extracellular space in an autophagy- and Cavin1-dependent process. Loss of Cavin1 expression could further exacerbate this process as by increasing cellular autophagy^64^. Autophagy-mediated secretion of Cav1 was specifically observed in LNCaP cells and could be a result of their stronger autophagic responses compared to other PCa cells including DU145 and PC3 cells^68^.

Autophagosomes form in the cytoplasm of cells and function primarily in the degradation of cellular proteins and organelles by the engulfment of unwanted cellular machinery within the maturing phagophore. Several autophagy-based pathways have been characterised to function in the non-canonical regulated release of proteins, which demonstrate a subversion of protein degradation for secretion into the extracellular space^47–51^. Cav1 degradation has been linked with autophagy^52^ as several studies have characterised specific interactions between Cav1 and known ATG family regulators^53, 54^. In agreement with these observations we demonstrated that inhibition of autophagy by siRNA knockdown of selected ATG proteins significantly increased cellular Cav1 levels following serum starvation for 6 hours (Fig. 7C-E). Additionally, treatment of cells with 3MA, a specific inhibitor of autophagosome maturation, resulted in a 3.2-fold increase in cellular Cav1 protein within treated cells following a 6-hour serum starvation (Fig. 7C-E). This was concurrent with a reduction in the release of Cav1 into the extracellular space (Fig. 7D and F). This model is entirely consistent with our observations and with the finding that even Cav1 tagged with a large GFP/YFP moiety is still efficiently released from cells.

The origin of the curvature of the particles released from LNCaP cells remains unknown. It is likely that the expression of caveolin itself is involved in sculpting the membrane to generate the released C-exosomes. We observed that Cav1 was the most abundant protein in C-exosomes based on our mass spectrometry analyses. In addition, we demonstrated that the S80E point mutant of Cav1, which disrupts caveola formation in mammalian cells and in a model prokaryotic system^34, 69^ potently inhibited secretion (Fig. 6). Moreover, this mutant was unable to efficiently form C-exosome precursors in the cytoplasm as judged by EM. The lipidic environment may also be a critical determinant for the formation of these vesicles in LNCaP cells. The S80A point mutant, which is known to increase cholesterol association^46^, resulted in increased Cav1 secretion compared to WT levels. Similarly, the S80E mutant which reduces cholesterol association^46^ inhibited the release of Cav1. Previous studies have shown that alterations to cellular cholesterol levels perturb the secretion of Cav1 from prostate cancer cells^24^ and over-expression of Apolipoprotein A-I preferentially induced the translocation of Cav1 and cholesterol onto small particles in the cytoplasm of rat astrocytes^66^. Other studies have suggested similarities between the properties of Cav1 particles released from LNCaP cells and high-density lipoprotein particles^19^. This suggests that cholesterol trafficking and Cav1 expression are tightly linked and cellular perturbation of cholesterol distribution may impact upon Cav1 secretion. In agreement with this, other studies have shown that the induction of Cavin1 expression in PCa cells resulted in widespread changes in the cellular distribution of cholesterol and correlated with reduced Cav1 secretion^15^. As putative precursor C-exosomes were observed in the cytoplasm of LNCaP cells but not PC3 cells, and this correlated with secretion of Cav1 in an inverted topology from LNCaP cells, this suggests differences in the way that these cells respond to Cav1. In view of the correlation between secreted lipids, particularly those characteristic of exosomes, and poor prognosis in prostate cancer^70^, the role of the lipid environment in Cav1 release warrants further investigation.

A molecular understanding of the pathway regulating C-exosome formation and secretion is a vital step in understanding the release of this potentially clinically relevant form of caveolin implicated in prostate cancer progression^12, 18, 60–62^. Studies into other proteins that are similarly released via secretory autophagy have not only demonstrated the importance of ATG genes (described here) but also ESCRT machinery, SNARE proteins and a dependence on Golgi reassembly stacking protein (GRASP) for secretion into the extracellular space^49, 50^. A detailed analysis of the proteins required for the regulated release of non-caveolar Cav1 will provide critical insights into prostate cancer, and potentially novel therapeutic targets to intervene for the inhibition of Cav1 secretion. We demonstrated Cavin1 expression inhibits the release of Cav1 from two separate PCa model cell lines. Other studies have shown the introduction of Cavin1 into a prostate cancer cell line in a tumour mouse model attenuated tumour progression^11^. This further suggests interesting avenues for therapeutic intervention and demonstrates the importance of understanding the molecular mechanisms underpinning Cav1 secretion.

## MATERIALS AND METHODS

### Tissue culture

Cells were grown in RPMI media (Gibco) supplemented with 10% Fetal Bovine serum and 2 mM L-glutamine. Cells were transfected using Lipofectamine 3000 (Invitrogen) as per the manufacturer’s instruction. siRNA knockdowns were performed as follows. LNCaP cells were seeded onto 35 mm dishes and left for 48 hours. siRNA oligos (Ambion, Life Technologies; siRNA ID #s s18158 and s18160) were transfected with Lipofectamine 3000 twice; 2nd and 3rd days after seeding. Cells were then transfected with YFP-Cav1, left for an additional 24 hours, and serum starved overnight.

### Antibodies and Reagents

Protease Inhibitor Cocktail Set III was purchased from Calbiochem. Ionomycin and 3MA were purchased from Sigma-Aldrich and Proteinase K from Roche. Protein A-Sepharose and Protein G-Sepharose were purchased from Sigma-Alderich. LDH release analysis was performed using CytoTox96 Non-radioactive Cytotoxicity Assay (Promega). Protein concentrations were assayed using BCA Protein Assay Kit (Quantum Scientific). Western blots were developed using Supersignal West Dura Extended Duration (Thermo Fisher Scientific). Released Cav1 was concentrated using Amicon Ultra-15 or 4, PLHK Ultracel-PL Membrane, 100 kDa (Merck). The following antibodies used in this study were raised in mouse unless otherwise stated; anti-Cav1 rabbit polyclonal antibody (BD Biosciences), rabbit anti-ATG5 (Sigma-Aldrich), rabbit anti-ATG9A (Abcam), rabbit anti-ATG12 (Abcam), rabbit anti-LC3B (Cell Signaling Technology), anti-CD63 monoclonal antibody (Developmental Studies Hybridoma Bank), anti-Myc-tag (Genesearch), anti-GFP/YFP (Roche Diagnostics Australia), anti-Actin (Millipore), anti-EEA1 (Becton Dickinson), anti-α-Tubulin (Sigma-Alderich) and anti-nucleoporin p62 raised in rabbit. Rabbit IgG was purchased from Sigma-Alderich. Alexa-555 secondary antibodies were purchased from Molecular Probes (Eugene, OR).

### Cav1 Isolation from conditioned media

Cells were grown until they reached 80% confluency then washed with phosphate buffered saline (PBS). Cells were then incubated for 16 to 48 h in phenol-free RPMI before harvesting. Exosome purification was performed by successive centrifugation of conditioned media. Media was centrifuged at 1500 x rpm for 5 min then 4500 x rpm for 20 min. LDH assays were performed on the media at this stage of centrifugation. Media was subsequently spun again 14,000 x *g* for 35 min to ensure all cells and cellular debris were removed. High-speed ultracentifugation was then performed for P100/S100 fractionation. Media was spun at 100,000 x *g* for 70 min using an Optima L-100XP floor standing ultracentrifuge (rotor SW40Ti). The supernatant was collected (S100 fraction) and concentrated using Amicon Ultra 100 kDa cut off concentrators. The pellet (P100 fraction) was resuspended in cold PBS, spun again at 100,000 x *g* for 70 min using an Optima MAX-XP bench-top ultracentrifuge (rotor TLS-55) and the pellet was collected. Whole cell lysates were analysed as follows. After conditioned media was removed, cells were washed in cold TNE buffer (100 mM NaCl, 0.1 mM EDTA and 50 mM Tris-HCl – pH 7.4) and scraped in the presence of 1% Triton-X100 and protease inhibitors. BCA assays were performed to determine protein concentration and LDH assay to determine percentage of cell death.

### Western Blots, Immunoprecipitation and sucrose gradients

Protein samples were boiled in sample buffer, separated using sodium dodecyl sulfate-polyacrylamide gel electrophoresis (SDS-PAGE; 10 – 15% acrylamide) and transferred onto PVDF membrane (Millipore). For CD63 western blots, protein samples were heated to 60°C for 10 min in a non-reducing buffer. Western blots were blocked in 3% bovine serum albumin in Tris Buffered Saline with Tween (TBST, pH 7.4 – 10mM Tris-HCl, 140mM NaCl and 0.1% v/v Tween 20), incubated with primary antibodies and HRP-conjugated secondary antibodies and developed using chemiluminescence. Immunoprecipitation was performed as follows. Supernatant was collected after the 14,000 x g spin and incubated with an anti-Cav1 antibody, anti-CD63 antibody or Rabbit IgG (as a negative control) for 1 h at 4°C in the presence or absence of 1% Triton-X100. 30 μL of Protein A-Sepharose beads were then added and incubated for 30 min. Beads were then centrifuged at 5000 x *g* for 2 min, and washed repeatedly in cold TNE buffer then processed as described above. Immunoprecipitation experiments using the P100 and S100 fraction were performed similarly but antibody incubation was performed after the second 100,000 x *g* spin. For sucrose gradients; samples were spun for 16 h at 55,000 rpm using Optima MAX-XP bench-top ultracentrifuge (rotor TLS-55).

### Proteinase K digestion

Purified P100 and S100 fractions were incubated with or without detergent at 4°C for 30 min on a shaker prior to ProK digestion. Proteins were then incubated with 250 ng/mL of ProK at 37°C for 30 min. Protease inhibitors were added after 30 min to stop the digestion.

### Fluorescence imaging

Coverslips were fixed in 4% paraformaldehyde in PBS at room temperature (RT) for 30 min, permeabilised with 0.1% saponin in PBS and quenched with 50 mM NH_4_Cl in PBS for 10 min. Coverslips were blocked in 0.2% bovine serum albumin and 0.2% fish skin gelatin in PBS for 10 min. Primary antibodies were incubated with coverslips for 1 h at RT, washed in PBS, and incubated with secondary fluorophores (Alexa555 anti-rabbit and Alexa660 anti-mouse) for 30 min. Coverslips were washed in PBS then water before mounting with Mowiol. Coverslips were imaged on a Zeiss LSM510 Confocal Microscope at 60X objective. Images for quantification were processed as follows. Individual GFP-Rab5Q79L endosomes were selected and the Red intensity was measured per endosome for multiple endosomes per cell for 30 to 50 cells per condition – repeated at least 3 times – such that an average red value can be compared between ± Cavin1. The area of GFP-Rab5Q79L endosomes was also compared to ensure no differences between the measured areas that existed between conditions.

### Proteomics

Isolated YFP-Cav1 particles and control particles were separated by SDS-PAGE to 8 mm. Staining, in-gel trypsin digest and LC-MS/MS were performed as previously described^71^. Spectrum Mill and Scaffold software were used for database searching and statistical analysis using normalized total precursor intensity. Report from Scaffold analysis is available as Table S1.

### Single molecule fluorescence spectroscopy

#### (i) Single molecule counting

The experiments were performed as described previously ^40^ using a commercial Zeiss 710 confocal microscope equipped with the ConfoCor module. Briefly, the 488nm excitation laser is focussed in the solution using a 40x water-immersion objective, creating a very small observation volume (∼1 fL). The fluorescence is collected, filtered using a 35-nm pinhole and recorded using single molecule counting detectors. The fluorescent Cav1 particles were diluted to picomolar concentrations to enable single particle detection. As described previously ^40, 41^, the diffusion of the Cav1 particles into the focal volume is recorded as a bright burst of fluorescence. The amplitude of the burst can be used to quantify the maximal number of proteins in the particles, after calibration of the brightness of YFP monomers.

#### (ii) Fluorescence Correlation Spectroscopy

FCS studies were performed exactly as described previously^40^. For these experiments, fluorescent proteins are diluted to 10-100 nanomolar concentration, so that a constant fluorescence intensity is detected. As fluorescent proteins enter or leave the detection volume constantly, the fluorescent intensity increases or decreases. The fluctuations of intensity around the average value are computed, and the auto-correlation of the intensity over time leads to a calculation of the diffusion time, the typical time it takes for a protein to diffuse through the focal volume. Binding between proteins or the formation of aggregates can be detected as the physical size, and consequently, the diffusion time, will increase upon complex formation.

### Negative Staining Electron Microscopy

Purified YFP-Cav1 particles were incubated for 10 min on glow-discharged carbon coated 1% formvar grids. Grids were washed 5 times in PBS, then 5 times in water and stained using 1% aqueous uranyl acetate. Purified exosomes from PC3 cells were adhered to formvar coated grids, washed repeatedly in PBS and water, then stained with 0.4% uranyl acetate in 2% methyl cellulose on ice for 10 min. Grids were imaged at 80 kV on a JEOL 1011 transmission electron microscope fitted with a Morada 4K X 4K Soft Imaging Camera at two-fold binning (Olympus).

### APEX-GBP Electron Microscopy

PC3, LNCaP and BHK cells were co-transfected with YFP or YFP-Cav1 and the APEX-GBP construct^42^. Cells were processed as described previously^42^. Briefly, cells were fixed in 2.5% glutaraldehyde, washed in 0.1M cacodylate buffer, and incubated with 3,3-diaminobenzoic acid (DAB – Sigma-Aldrich) in the presence of hydrogen peroxide for 30 min at room temperature. Cells were washed in cacodylate buffer, post-fixed in 1% osmium tetroxide for 2 min, serially dehydrated in ethanol and serially infiltrated with LX112 resin. Resin was polymerised at 60°C overnight and 60 nm ultrathin sections were cut on an Ultracut 6 (Leica) ultramicrotome and imaged as described above.

## ACKNOWLEDGEMENTS

The authors would like to thank Prof. T Thompson for sharing reagents, Dr. Vikas Tillu and Ms Dorothy Loo for their technical assistance on this project, and Ye-Wheen Lim and Natasha Kaushik for proofreading the manuscript. The authors acknowledge the use of the Australian Microscopy and Microanalysis Research Facility at the Center for Microscopy and Microanalysis at The University of Queensland. Fluorescence Microscopy was performed at the Australian Cancer Research Foundation (ACRF)/Institute for Molecular Bioscience (IMB) Dynamic Imaging Facility for Cancer Biology. Mass Spectrometry was performed at The University of Queensland Diamantina Institute Mass Spectrometry Facility. This work was supported by grants and a fellowship from the National Health and Medical Research Council of Australia (grant numbers APP1058565 and APP569542 to RGP; APP1045092 to RGP and NA; APP1037320 to RGP and KA; APP1108859 to YG and NA). YG and MMH were supported by separate ARC Future Fellowships (FT110100478 and FT120100251) and FAM with an ARC Discovery Project (DP170100125). RDT and FAM are supported by separate NHMRC Research Fellowships (APP1041929 and APP1060075).

## AUTHOR CONTRIBUTIONS

N.A. designed experiments, performed electron microscopy, biochemistry, and wrote and edited the manuscript. Y.W designed experiments, performed biochemistry and fluorescence microscopy, and wrote and edited the manuscript. S.O. designed experiments, performed biochemistry and fluorescence microscopy and interpreted data. Y.G. performed the FCS and single molecule fluorescence spectroscopy. J.F. performed confocal microscopy. C.F. and J.R. provided technical expertise for electron microscopy. M.M.H. designed the proteomic analyses, analysed and interpreted data and edited the manuscript. F.A.M., K.A. and R.D.T. interpreted data. R.G.P. designed experiments, interpreted data, edited the manuscript, and supervised the work. All authors reviewed the results and approved the final manuscript.

## Supplementary Data

**Figure S1.**
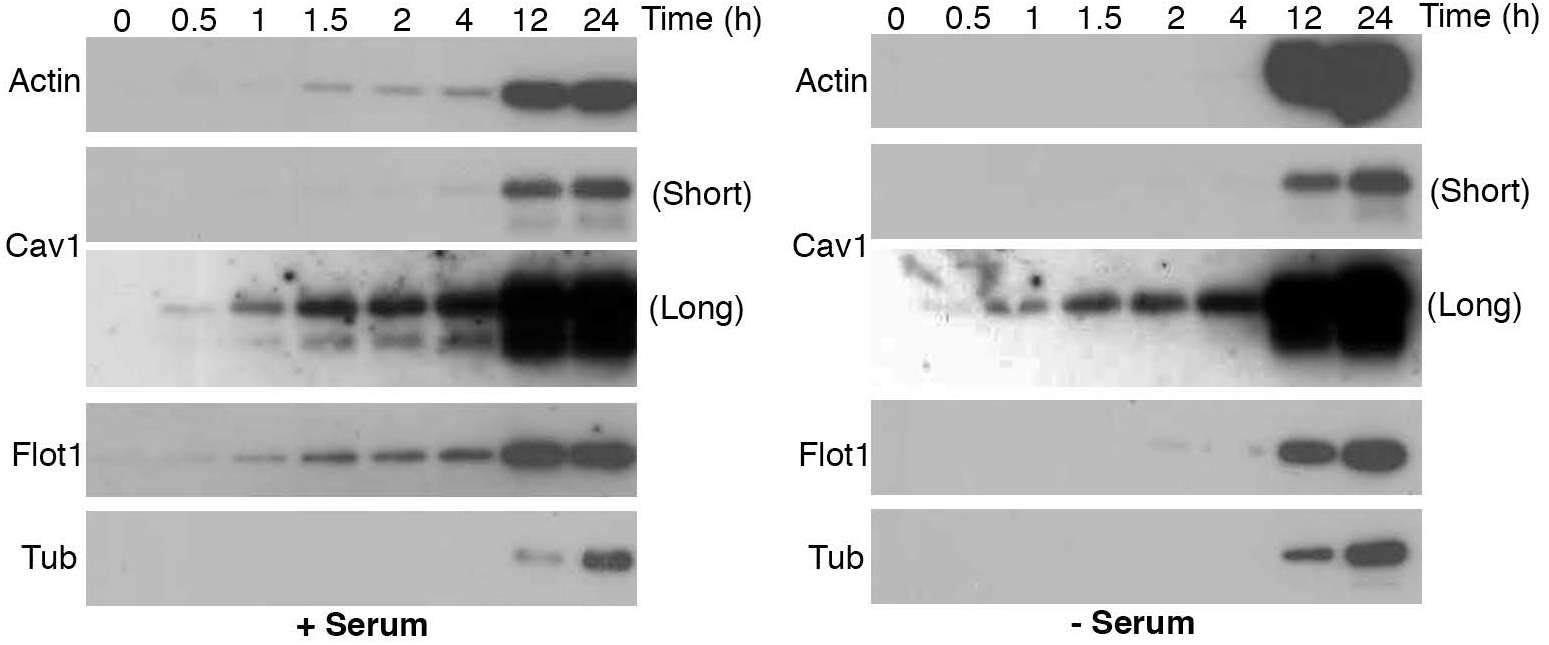
Time course of exosomal secretion from PC3 cells. Cav1, Flot1, actin and tubulin are secreted with or without serum indicating Cav1 detection in the media is a direct result of release from cells and not a consequence of Cav1 presence in serum exosomes. Moreover, independence from starvation suggests autophagy is not required for Cav1 secretion in PC3 cells.

**Figure S2.**
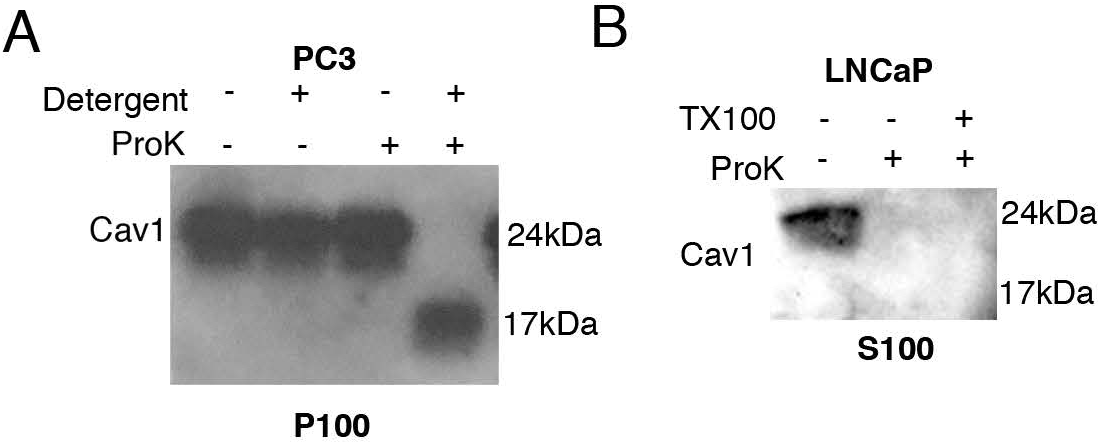
A) Western blot of ProK-treated PC3 P100 fraction demonstrating Cav1 is unavailable for digestion unless membranes are disrupted with β-octylglucoside and Triton-X100. B) Proteinase-K mediated degradation of Cav1 isolated from the LNCaP S100 fraction results in a complete loss of the α-Cav1 recognition epitope without TX100 detergent treatment.)

**Figure S3.**
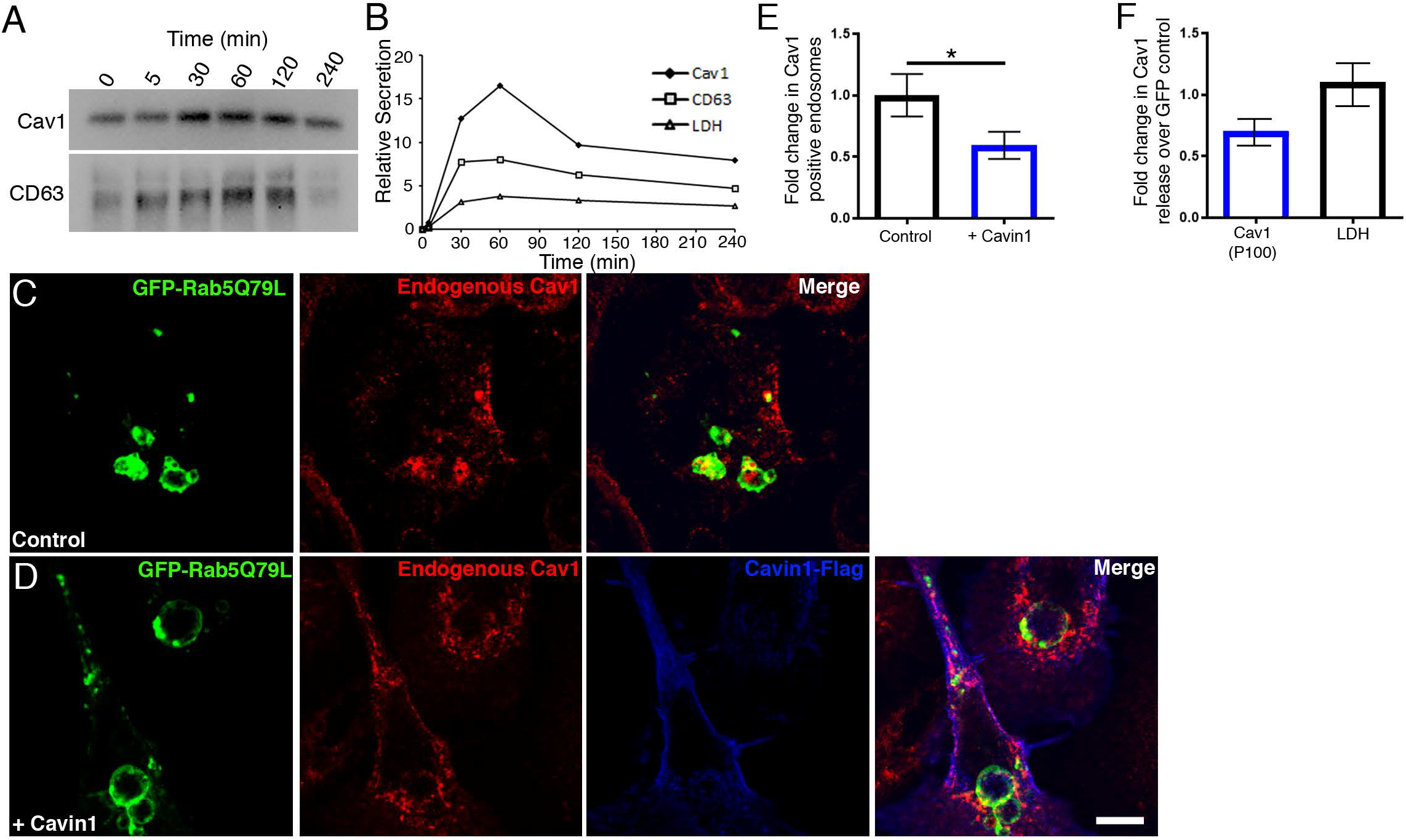
A) Time course of Cav1 release after ionomycin treatment. Cells were treated with serum free media or 500 nM ionomycin. B) Peak secretion was observed at 30 to 60 minutes of treatment. C) PC3 cells transiently transfected with GFP-Rab5Q79L accumulate Cav1 within enlarged endosomes. D) PC3 cells co-transfected with Cavin1 reduces the sorting of Cav1 into GFP-Rab5Q79L positive endosomes. E) Two-tailed t-test demonstrating a significant reduction in Cav1-positive endosomes when Cavin1 is expressed. p = 0.0418, n = 4 with each independent repeat analyzing > 20 cells per experimental condition, error bars represent SEM. F) Fold change in of Cav1 release from PC3 cells stably expressing GFP-Cavin1 compared to GFP. n = 3, error bars represent SEM.

**Table S1. Proteomics analysis of YFP-Cav1 particles.** Proteome composition of GFP-trap pulldowns performed on conditioned media collected YFP-Cav1 or YFP expressing LNCaP cells (n=3) were analysed by SDS-PAGE gel separation, in-gel digest and LC-MS/MS analysis. Proteins that were significantly different (p<0.05) between YFP-Cav1 and YFP groups across the three replicates are highlighted in red font. Parameters and results from Scaffold analysis are shown in this document.

